# Targeting the acetyltransferase NAT10 corrects pathologies in human frontotemporal dementia neurons and extends lifespan in an in vivo *Drosophila* tauopathy model

**DOI:** 10.1101/2024.10.19.619211

**Authors:** Francesco Paonessa, Bernardo Delarue Bizzini, Tom Campbell, Ravi Solanki, Richard Butler, James Smith, Catherine M. Davidson, Delphine Larrieu, Andrea H. Brand, Frederick J. Livesey

## Abstract

Disruption of the neuronal nuclear membrane and perturbation of nucleocytoplasmic transport are features of neurodegenerative diseases, including Alzheimer’s disease, that involve the microtubule-associated protein tau (MAPT). We previously identified that missense and splicing mutations in the MAPT gene, causal for frontotemporal dementia, result in nuclear envelope deformation and disrupted nucleocytoplasmic transport in human neurons. This is most likely due to microtubule mechanical stress, similar to that observed in Hutchinson-Gilford Progeria Syndrome (HGPS). A small molecule inhibitor of the acetyltransferase NAT10 has been shown to correct nuclear membrane defects in HGPS by modulating microtubule dynamics. We report here that NAT10 inhibition alters microtubule dynamics and corrects nuclear lamina defects and aberrant nucleocytoplasmic transport in human iPSC-derived FTD-MAPT neurons. Similarly, NAT10 inhibition and haploinsufficiency correct neuronal nuclear shape defects and extend lifespan *in vivo* in a *Drosophila* model of tauopathy. Our results show that NAT10 mediates neuronal pathologies in tauopathies and is a promising new therapeutic target in these diseases.

## INTRODUCTION

The microtubule-associated protein tau (MAPT) is central to the pathogenesis of many neurodegenerative diseases, forming intracellular aggregates or neurofibrillary tangles in diseases collectively referred to as tauopathies, including Alzheimer’s disease, progressive nuclear palsy and frontotemporal dementia (Creekmore et al., 2024; Goedert et al., 2024). Autosomal dominant mutations in the gene encoding MAPT are causal for a highly penetrant form of frontotemporal dementia (FTLD-tau), with more than 50 missense and intronic MAPT mutations identified as causal for FTD (Creekmore et al., 2024; Ghetti et al., 2015). The majority of missense mutations are in and around the microtubule-binding region, promoting protein aggregation, whereas the intronic mutations alter splicing of the repeat-encoding exon 10, increasing the amount of 4-microtubule binding repeat (4R) form relative to the 3-repeat (3R) form (Creekmore et al., 2024).

Previously, we reported that both intronic (MAPT IVS10+16) and missense (MAPT P301L) MAPT mutations are associated with abnormalities of the nuclear membrane in human neurons *in vitro* and *in vivo*, with disrupted nucleocytoplasmic transport *in vitro* (Paonessa et al., 2019). Similar tau-associated nuclear shape abnormalities have been reported in Huntington’s disease (Fernández-Nogales et al., 2014) and Alzheimer’s disease (Frost et al., 2014). Tau-mediated disruption of the nuclear pore complex has been proposed as the pathogenic mechanism altering nucleocytoplasmic transport in Alzheimer’s disease (Eftekharzadeh et al., 2018). *Drosophila* models of tauopathy also exhibit nuclear abnormalities, with accumulation of mRNA in and around nuclear invaginations (Cornelison et al., 2019; Frost et al., 2016), together with disrupted heterochromatin (Frost et al., 2014).

The occurrence and effects of the neuronal nuclear abnormalities observed in AD and FTD – nuclear lamina disruption, alterations in nucleocytoplasmic transport, and changes in global heterochromatin – are similar to those identified in Hutchison-Gilford Progeria Syndrome (HGPS), a rare inherited laminopathy characterized by accelerated aging of non-neural tissues (Pollex and Hegele, 2004). HGPS results from a mutation in the *LMNA* gene that impairs the correct processing of prelamin A protein (Pollex and Hegele, 2004). This leads to the accumulation of a shorter and toxic form of lamin A, known as progerin, causing disruption of nuclear membrane integrity and a characteristic cellular phenotype of nuclear blebbing. In cellular models of HGPS there is a disruption of cytoskeletal and microtubule dynamics as well as alterations in nucleocytoplasmic transport (Balmus et al., 2018; Larrieu et al., 2018, 2014). In HGPS cell models, we found that the small molecule Remodelin, an inhibitor of the NAT10 acetyltransferase, corrects both nuclear shape abnormalities and the disruption of nucleocytoplasmic transport (Larrieu et al., 2014). Furthermore, both Remodelin treatment and haploinsufficiency of NAT10 increased health and lifespan in a mouse model of HGPS (Balmus et al., 2018).

Given the similarities in the cellular phenotypes in human FTD-MAPT neurons and HGPS fibroblasts, we investigated the effect of inhibiting NAT10 in human tauopathy models *in vitro* and in a *Drosophila* tauopathy model *in vivo*. We find that inhibition of NAT10 corrects nuclear invaginations in human iPSC-derived FTD-MAPT neurons and reduces microtubule elongation and mechanical stress on the nuclear envelope, improving nucleocytoplasmic transport. Furthermore, knock-down of NAT10 also corrects nuclear lamina defects in FTD neurons. In a *Drosophila* tauopathy model, inhibition of dNAT10 or haploinsufficiency of dNAT10 corrected nuclear membrane abnormalities and significantly extended animal lifespan. These results indicate that NAT10 mediates microtubule-mediated nuclear pathologies in FTD neurons and may represent a promising therapeutic target for tauopathies.

## RESULTS

### A small molecule inhibitor of the NAT10 acetyltransferase corrects nuclear lamina defects in human FDT-MAPT neurons

In a previous study, we found that pathogenic mutations in the MAPT gene, causal for frontotemporal dementia, resulted in abnormal microtubule dynamics in the neuronal cell body, deforming the nuclear membrane and disrupting nucleocytoplasmic transport (Paonessa et al., 2019). These phenotypes are very similar to those observed in non-neural cells in the premature ageing condition, Hutchinson-Gilford Progeria Syndrome (HGPS), which are corrected by a small molecule inhibitor of the NAT10 acetyltransferase, Remodelin (Larrieu et al., 2014). Therefore, we first investigated whether nuclear lamina defects in human FTD-MAPT neurons are affected by NAT10 inhibition.

To do so, cortical excitatory neurons were generated from iPSCs derived from two different individuals carrying the pathological MAPT IVS10+16 mutation (MAPT IVS10+16-A and MAPT IVS10+16-B), an isogenic corrected line (MAPT IVS10+16 B-isogenic) and an unrelated, independent non-demented control line. Ninety (90) days from the initiation of neuronal differentiation, neurons were treated with the NAT10 inhibitor Remodelin for 48 hours and the frequency of nuclear lamina invaginations in neurons scored based on Lamin B1 staining (see Methods for details).

Consistent with our previous data, FTD-MAPT neurons had a higher frequency of nuclear invaginations compared to control neurons (Figure 1A, B). Remodelin treatment reversed this phenotype, reducing the proportion of neurons with nuclear lamina defects to control levels (Figure 1A, B).

**Figure 1.**
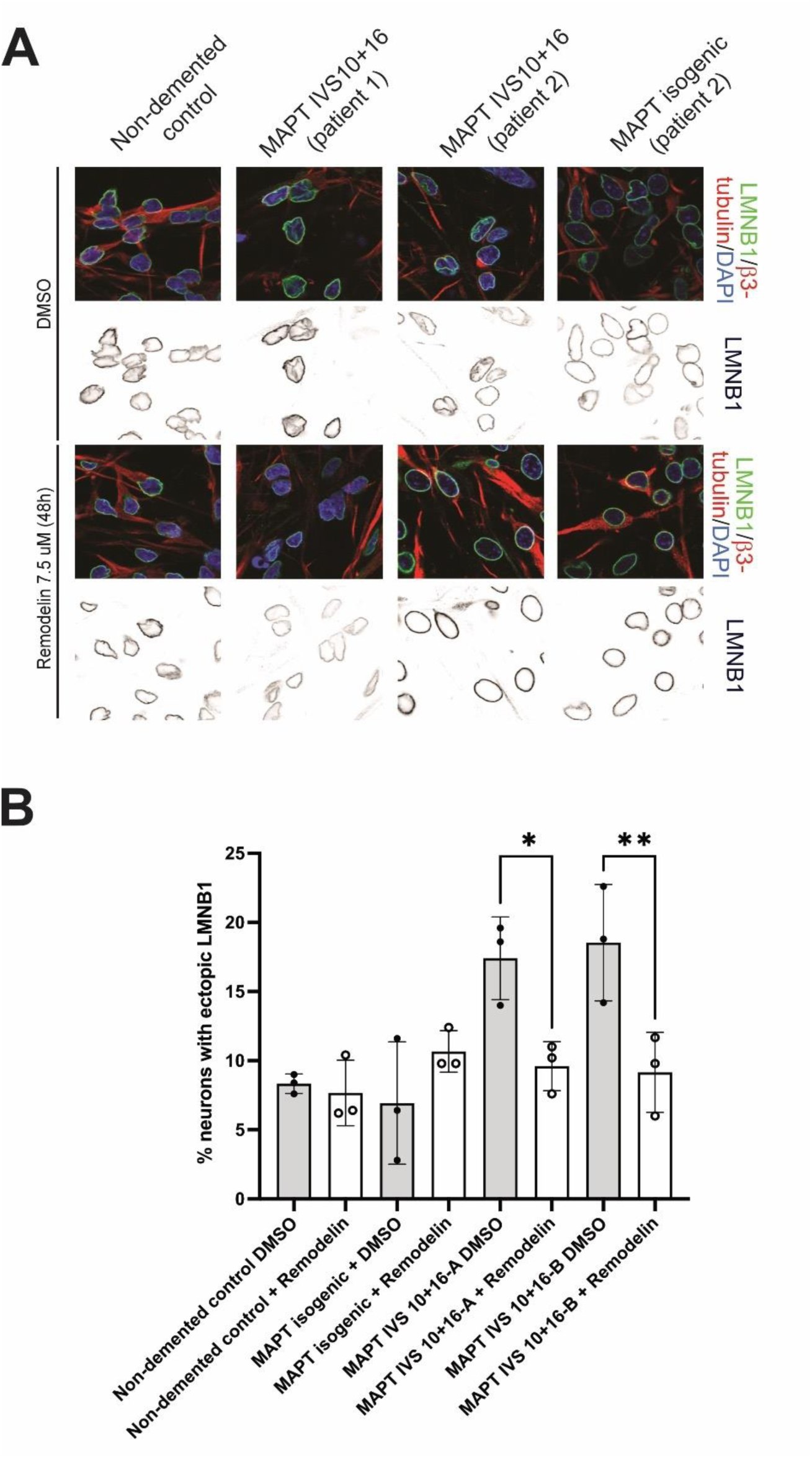
The NAT10 small molecule inhibitor Remodelin corrects nuclear lamina defects in human FTD-MAPT neurons. **A.** Confocal images of the nuclear lamina (lamin B1, green) in FTD-MAPT neurons (MAPT IVS10+16-A and MAPT IVS10+16-B) compared with non-demented and MAPT IVS10+16-B isogenic control neurons (120 DIV) (β3-tubulin, red; DAPI, blue). Neurons were exposed to 7.5μM Remodelin or vehicle control (DMSO) for 48 hours. Abnormalities of nuclear lamina shape are common in FTD-MAPT neurons, while nuclear lamina invaginations are reduced by inhibition of NAT10 with Remodelin. **B.** FTD-MAPT neurons had an increased incidence of neurons with nuclear lamina invaginations compared with control neurons (grey bars) in DMSO, which was significantly reduced after treatment with Remodelin (white bars). Significance was determined by one-way ANOVA followed by Tukey’s test (*p < 0.05; **p < 0.01); error bar represents SD; n = 3 independent experiments, dots represent the average of one experiment).

### Small molecule inhibition of NAT10 corrects neuronal nuclear defects and extends survival in a *Drosophila* tauopathy model

*Drosophila melanogaster* is a well-established system for in vivo modelling of a number of neurodegenerative diseases, including tauopathies (McGurk et al., 2015). Transgenic expression of human MAPT variants encoding missense mutations casual for FTD capture several features of disease, including neuronal degeneration, together with abnormal movement and reduced lifespan (Frost et al., 2016, 2014; Wittmann et al., 2001). Disruption of the nuclear lamina and chromatin reorganization have previously been reported in adults expressing pan-neuronal human MAPT R406W (Frost et al., 2016), which also shortened their lifespan (Wittmann et al., 2001).

To determine whether inhibition of NAT10 with Remodelin corrects nuclear membrane defects *in vivo*, MAPT R406W-expressing animals were maintained on food containing either vehicle or 100μM Remodelin, the highest concentration without developmental toxicity (Supplementary Figure 1). As previously reported (Frost et al., 2016), pan-neuronal expression of MAPT R406W resulted in neuronal nuclear lamina invaginations in adults (Figure 2A, B). Remodelin treatment reduced the number of neuronal nuclei with nuclear membrane defects in the *Drosophila* tauopathy model compared to the control group (Figure 2A, B), consistent with the effect of NAT10 inhibition in human FTD-MAPT neurons.

**Figure 2.**
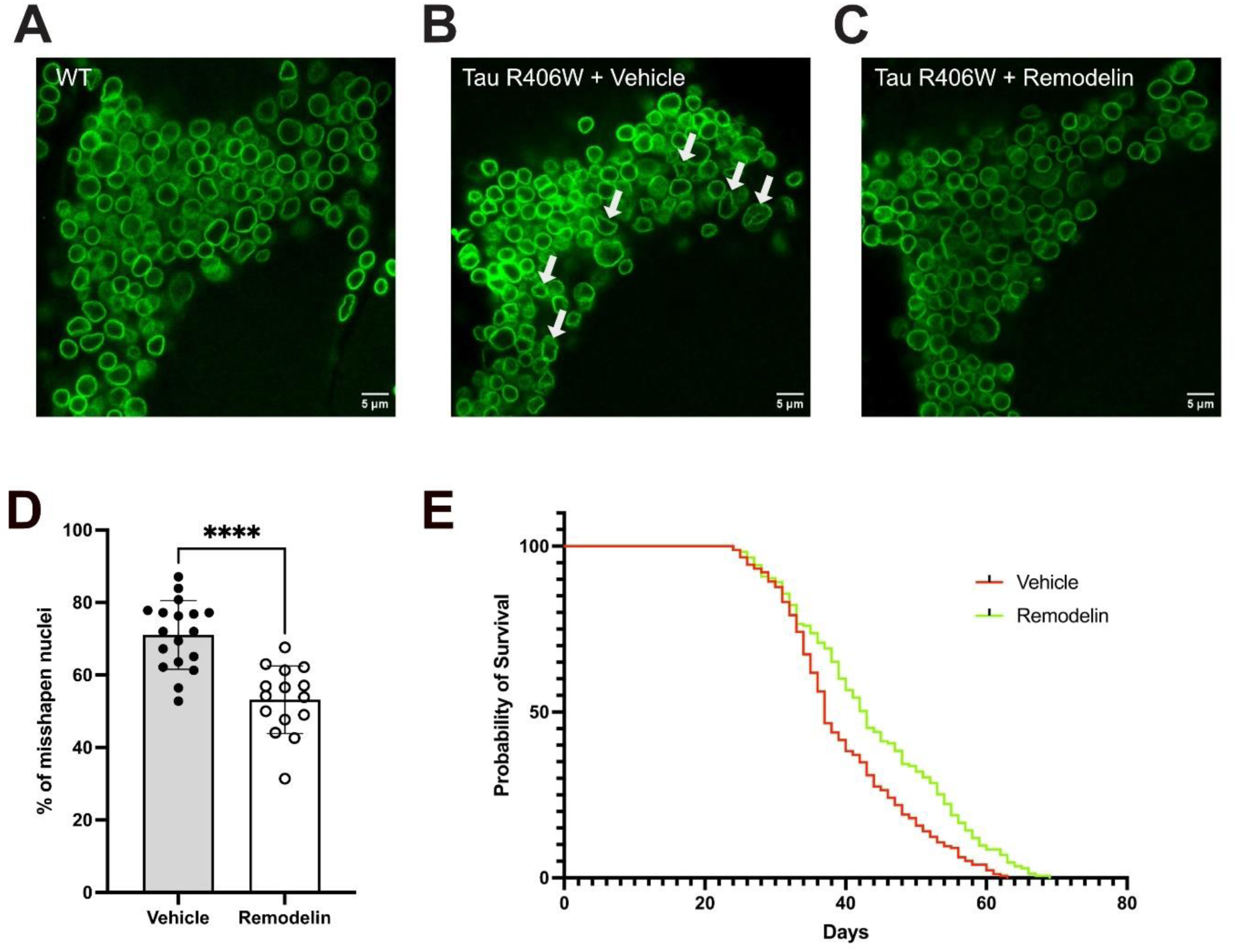
The NAT10 small molecule inhibitor Remodelin corrects nuclear lamina morphology defects and extends lifespan in the Drosophila MAPT R406W tauopathy model. **A-C.** Confocal images of the nuclear lamina (Lamin B, green) in neurons on the ventral side of the antennal lobe of 44-day-old control (**A**; w1118 ; + ; +) and elaV > MAPT R406W *Drosophila*, fed on either vehicle (**B**) or Remodelin-containing food (**C)**. Animals pan-neuronally expressing MAPT R406W were exposed to food containing either vehicle (20% DMSO, 65% 2-Hydroxypropyl-b-cyclodextrin and 15% Tween 80) or 100 μM Remodelin throughout development and lifespan. Neurons in the antennal lobe displayed frequent nuclear lamina defects in vehicle-treated MAPT R406W (arrows), whereas neurons from animals exposed to Remodelin show improved nuclear morphology. Scale bars, 5 μm **D.** Quantification of the misshapen nuclear frequency was performed on 44-day-old *Drosophila* neurons expressing elaV > MAPT R406W. Animals exposed to vehicle alone show an increased percentage of misshapen neuronal nuclei compared to neurons from animals of the same age treated with Remodelin. Significance was tested by unpaired Student’s t-test (p**** < 0.0001); error bar represents SD; n = 18 (vehicle) and n = 15 (Remodelin) representing one section per antennal lobe and hemisphere (max. 2 sections per animal), whereas dots represent the percentage of misshapen nuclei per brain area scored. **E.** A lifespan experiment was performed to assess the effect of Remodelin on longevity of *Drosophila* pan-neuronally expressing MAPT R406W. Remodelin treatment throughout development and adult life increased median and total lifespan compared to the vehicle-only group. Significance was determined using the log-rank test (p**** < 0.0001); Survival was recorded for n = 178 (Vehicle) and n = 175 (Remodelin) female flies.

Transgenic expression of both wild-type and mutant forms of human tau are associated with shortened lifespan in *Drosophila* (Gistelinck et al., 2012).Therefore, we investigated whether NAT10 inhibition affected overall fitness in transgenic animals expressing human MAPT R406W. We found that Remodelin exposure increased overall and median survival when compared to animals exposed to vehicle alone (Figure 2C). Thus, small molecule inhibition of NAT10 both prevented tau-mediated neuronal nuclear membrane defects *in vivo* and extended lifespan in animals expressing the pathogenic MAPT R406W variant.

### NAT10 inhibition alters microtubule dynamics in human FTD-MAPT neurons

Deformation of nuclear membranes in FTD-MAPT neurons is associated with altered microtubule dynamics within the neuronal cell body, with microtubules deforming the nucleus (Paonessa et al., 2019). Acute treatment with the microtubule-destabilizing compound, nocodazole, restored nuclear shape, demonstrating that nuclear membrane deformation is a dynamic process (Paonessa et al., 2019). Therefore, we investigated whether NAT10 inhibition with Remodelin impacts microtubule dynamics in both control and FTD-MAPT neurons.

To do so, we introduced a construct driving expression of the microtubule tip-binding protein, EB3, fused to GFP and driven by the neuron-specific promoter, Synapsin1 (*pSyn1*:EB3-GFP) into control and FTD-MAPT human neurons (Figure 3). Neuronal expression of GFP was observed within one day, and most cells were positive within three days. Single-plane videos of EB3-GFP were captured enabling microtubule growth to be tracked and analyzed in individual neurons (Figure 3 and Supplementary Movies 1-4).

**Figure 3.**
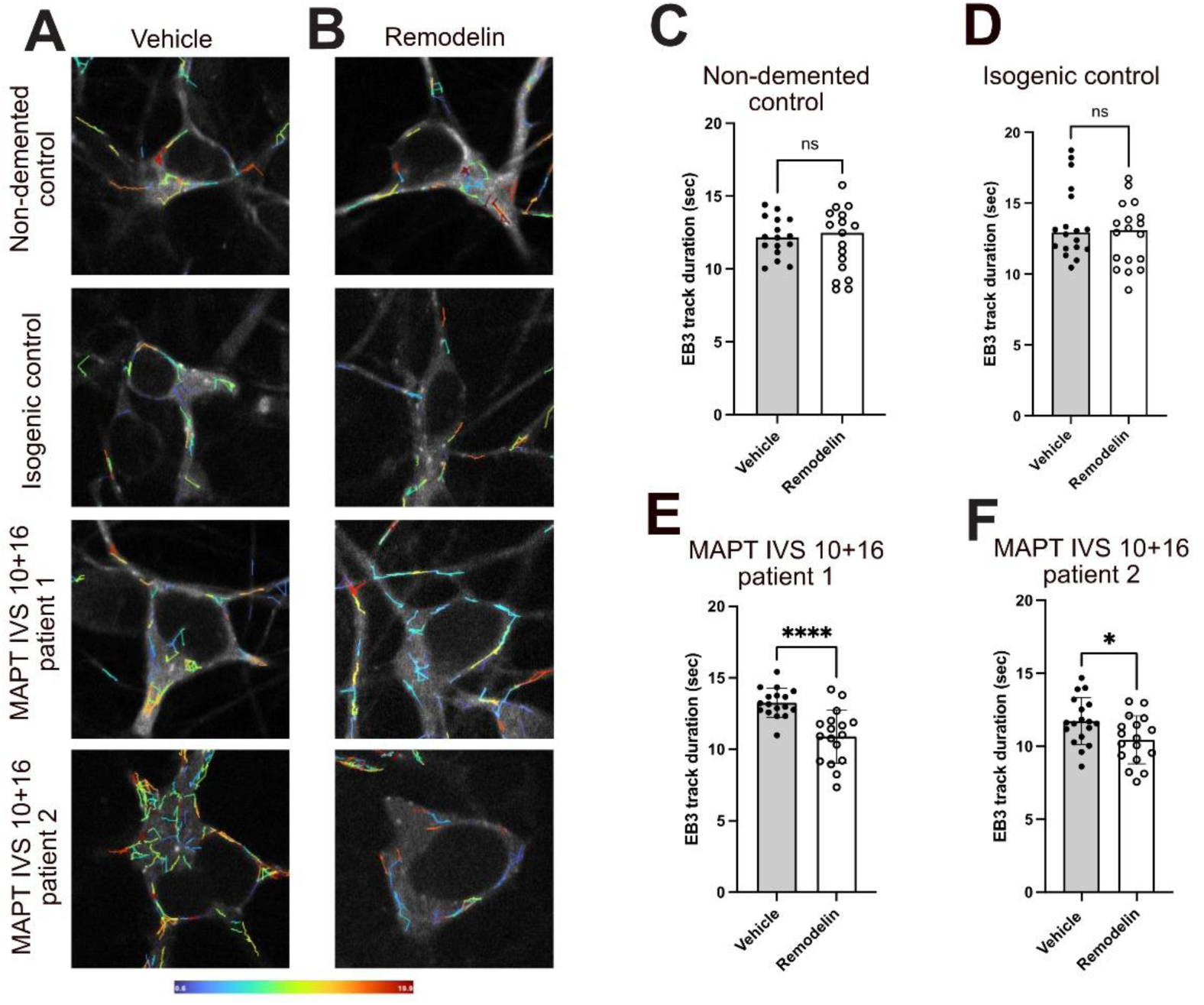
NAT10 inhibition alters microtubule dynamics in human FTD-MAPT neurons. **A-B.** Representative images of microtubule track duration (cumulative over a 150 sec interval movie), visualised by live-imaging of an EB3-GFP fusion protein, overlaid on stills from GFP-EB3 live imaging (gray) of control and FTD-MAPT-neurons (120 DIV) after 24 hours treatment with Remodelin or vehicle (DMSO). **C-F.** Microtubule growth, reflected in EB3 track duration, is reduced by Remodelin treatment in FTD-neurons but not in control neurons. Significance was determined using Student’s t test (ns = not significant; *p < 0.05; ****p < 0.0001); error bar represents SD; n > 15 movies from 3 independent experiments, dots represent the average of one movie).

FTD neurons exposed to Remodelin for 24 hours demonstrated a reduction in EB3-GFP track duration, whereas Remodelin had no effect on EB3 track duration in isogenic and unrelated non-demented control neurons (Figure 3). EB3-GFP track duration is a measure of microtubule growth. A reduction in track duration indicates an overall reduction in microtubule elongation within FTD neurons following NAT10 inhibition. This decrease in microtubule elongation, which accompanies the correction of nuclear lamina defects, is consistent with our previous observation that nocodazole treatment rectifies nuclear lamina defects (Paonessa et al., 2019). Therefore, it is likely that NAT10 inhibition corrects nuclear lamina defects by modulating microtubule dynamics, reducing mechanical stress on the nuclear membranes of FTD neurons.

### Nucleocytoplasmic transport defects in FTD-MAPT neurons are reversed by inhibition of NAT10

Impaired nuclear membrane integrity and disrupted nucleocytoplasmic transport are significant pathologies in several neurodegenerative diseases, including ALS and ALS-FTD (Kim and Taylor, 2017). More recently, disruption of nuclear pore complexes and aberrant nucleocytoplasmic transport have been found in ALS due to *C9ORF72* expansion (Zhang et al., 2015), in Alzheimer’s disease (Eftekharzadeh et al., 2018) and FTD-MAPT (Paonessa et al., 2019). In the latter study, we found that the aberrant nucleocytoplasmic transport in human FTD-MAPT neurons was corrected by acute exposure to the microtubule depolymerizing small molecule nocodazole (Paonessa et al., 2019).

Therefore, we tested whether the correction of nuclear lamina defects by NAT10 inhibition and the reported effect on microtubules was accompanied by an improvement in nucleocytoplasmic transport. To do so, we carried out fluorescent recovery after photobleaching (FRAP) experiments in iPSC-derived neurons expressing Shuttle-tdTomato, a NLS-NES-tagged TdTomato (Zhang et al., 2015) that dynamically moves between the nucleus and cytoplasm. As expected, Shuttle-tdTomato was enriched in the nucleus of infected iPSC-derived neurons, with a less in the cytoplasm and neurites (Figure 4A). Our working hypothesis was that increased nucleus-cytoplasm diffusion in FTD-MAPT neurons that we previously reported (Paonessa et al., 2019) would result in a faster recovery rate of tdTomato signal after photobleaching within the nucleus, compared to healthy control neurons.

**Figure 4.**
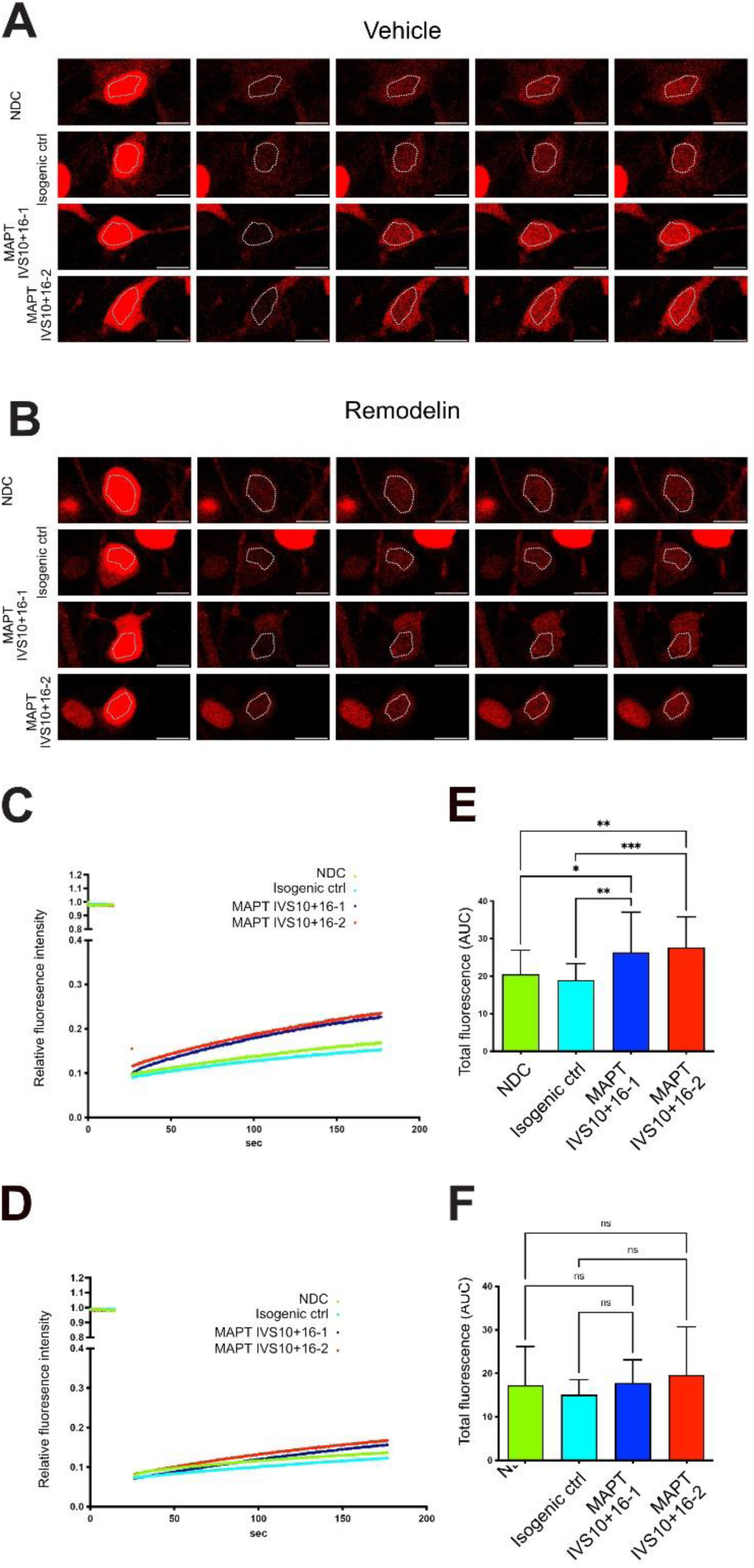
Nucleocytoplasmic transport defects in FTD-MAPT neurons are reversed by inhibition of NAT10. Control and FTD-neurons (120 DIV) were transduced with lentivirus expressing a NLS:TdTomato:NES reporter and subjected to <u>F</u>luorescence <u>R</u>ecovery <u>A</u>fter <u>P</u>hotobleaching (FRAP) after 24h treatment with DMSO (vehicle) or Remodelin. **A-B.** Representative images of neurons treated with DMSO (vehicle) or Remodelin before bleaching of nuclear TdTomato and then at t=0 sec and t=150 sec post-bleaching. **C-D.** Relative TdTomato fluorescence intensity within the nucleus over a period of 180 sec following photobleaching. Vehicle-treated FTD-neurons show a quicker recovery of nuclear FRAP of TdTomato relative to non-demented control neurons (C). After treatment with Remodelin, nuclear TdTomato shows a similar recovery rate between control and FTD-neurons following photobleaching (D). Curves represent the average of > 15 movies. **E-F.** Area Under the Curve (AUC) of total nuclear FRAP of TdTomato from individual movies. Significance was determined using one-way ANOVA followed by Tukey’s test (ns = not significant; *p < 0.05; **p < 0.01; ***p < 0.001); error bar represents SD; n > 15 movies from 3 independent experiments). MAPT-FTD neurons demonstrate statistically significantly more nuclear FRAP compared to non-demented control neurons when treated with vehicle. Following Remodelin treatment, nuclear FRAP in MAPT-FTD neurons is indistinguishable from that of non-demented controls.

To test whether this is the case, and if it could be corrected by inhibition of NAT10, neurons expressing Shuttle-tdTomato were treated with vehicle (DMSO) or Remodelin for 24 hours before FRAP analysis in neuronal nuclei. Vehicle-treated FTD-MAPT neurons showed a faster recovery of nuclear tdTomato signal compared to either NDC or MAPT IVS10+16B isogenic control neurons, indicating aberrant nucleocytoplasmic transport (Figure 4A, C and E), consistent with our previous study (Paonessa et al., 2019). Inhibition of NAT10 by treatment with Remodelin for 24 hours slowed the recovery of nuclear tdTomato in FTD-MAPT neurons, such that the rate of FRAP was indistinguishable between NAT10-treated FTD-MAPT, isogenic and unrelated NDC control neurons (Figure 4B, D and F). Therefore, NAT10 inhibition with Remodelin corrects nucleocytoplasmic transport defects in MAPT-FTD neurons.

### Reduction of NAT10 protein corrects nuclear lamina defects in human FTD-MAPT neurons

To confirm that the effects of Remodelin in human neurons are via its known target, NAT10, we sought to generate NAT10 loss of function FTD-neurons. Consistent with evidence that NAT10 is an essential gene for development in mice (Balmus et al., 2018), we could not successfully generate NAT10 knockout human iPSCs by CRISPR-mediated gene targeting (data not shown).

As an alternative strategy, we acutely knocked down NAT10 in neurons using siRNAs (NAT10 KD). Following siRNA delivery, NAT10 mRNA and protein were both reduced by over 60% in neurons, without impacting neuronal viability (Figure 5 A-C). NAT10 KD corrected the frequency of nuclear lamina defects in FTD-MAPT neurons compared to a scrambled control siRNA (Figure 5), and to a comparable degree as Remodelin inhibition of NAT10 (Figure 5). Therefore, acute loss of NAT10 function, or small molecule inhibition of its acetyltransferase activity, corrects nuclear lamina defects in human FTD-MAPT neurons.

**Figure 5.**
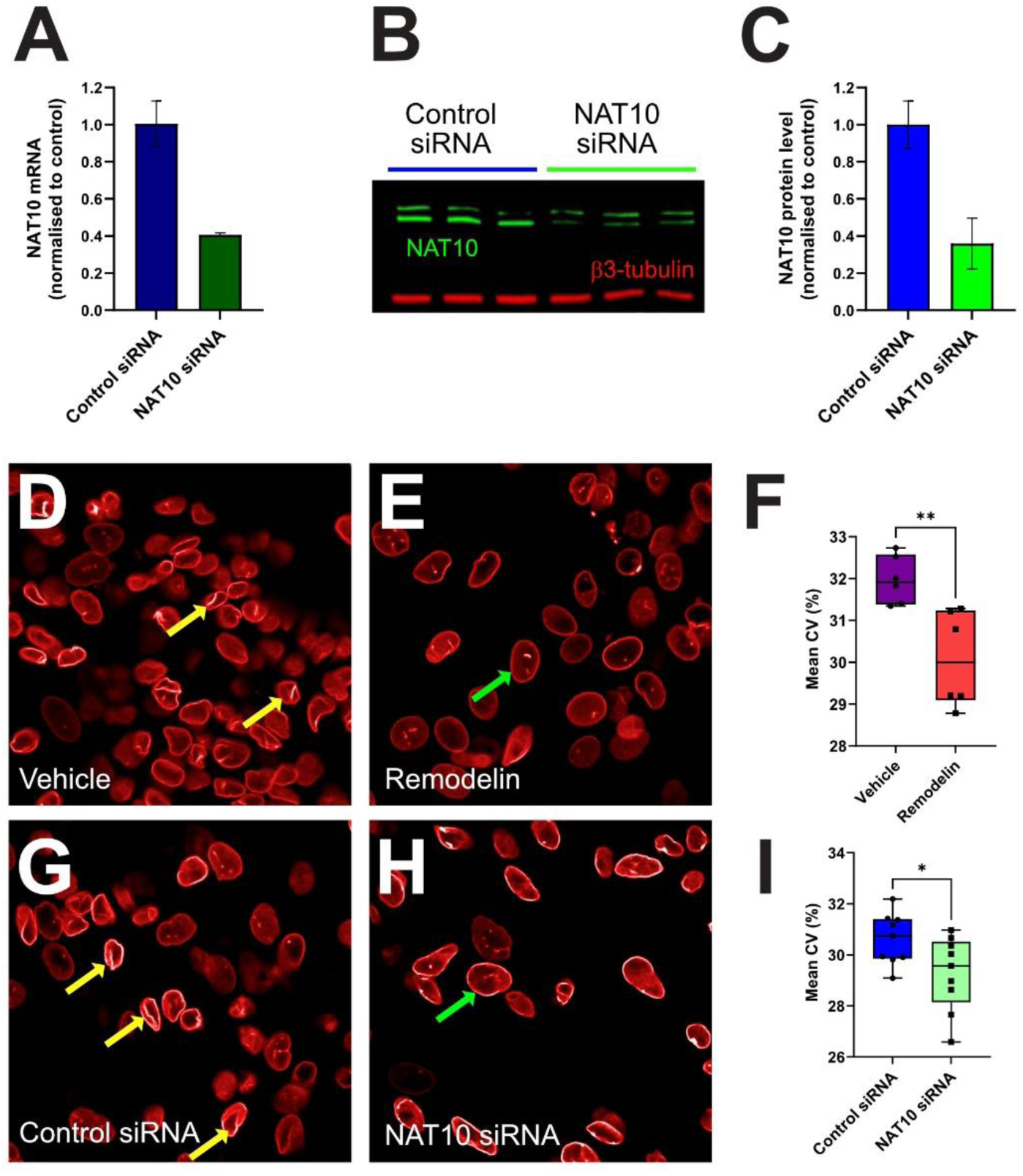
Reduction of NAT10 protein corrects nuclear lamina defects in human FTD-MAPT neurons. NAT10 mRNA was acutely knockdown of in human MAPT-FTD neurons by treatment with NAT10-targeting or control siRNAs for 72 hours. **A.** NAT10 siRNA reduces NAT10 mRNA by over 60%, compared to control siRNA treatment. **B-C.** NAT10 siRNA treatment for 72 hours reduces NAT10 protein by over 60%, compared with control siRNA treatment. **D-F.** Remodelin treatment significantly reduces the incidence of nuclear lamina defects in MAPT-FTD neurons, compared with vehicle-treated neurons (DMSO). Yellow arrows indicate nuclei with nuclear lamina defects, green arrows indicate examples of normally-shaped nuclei. Significance was determined by Student’s t test (*p < 0.05; **p < 0.01); error bar represents SD). **G-I.** Reduction of NAT10 protein by siRNA knockdown significantly reduces the incidence of nuclear lamina defects in MAPT-FTD neurons, compared with control siRNA-treated neurons. Yellow arrows indicate nuclei with nuclear lamina defects, green arrows indicate examples of normally-shaped nuclei. Significance was determined using Student’s t test (*p < 0.05; **p < 0.01); error bar represents SD).

### *Drosophila* NAT10 haploinsufficiency corrects nuclear defects and extends lifespan in an *in vivo* tauopathy

We tested whether reducing NAT10 *in vivo* corrects tauopathy phenotypes in the *Drosophila* MAPT-R406W model. The *Drosophila* orthologue of NAT10, l(1)G0020 (dNAT10) shares 53% homology with human NAT10, and the acetyltransferase domain of dNAT10, the likely target of Remodelin, is conserved between the species. The dNAT10 mutant line (l(1)G0020(1)/FM7c ; + : +; Bloomington Drosophila Stock Center #11474) is homozygous lethal: only balanced males (FM7c/y) and heterozygous or homozygous balanced females (l(1)G0020/FM7c or FM7c/FM7c) emerge as adults. This is consistent with our results that NAT10 KO iPSCs are not viable, and with the previously reported lethality of homozygous NAT10 mutant mice (Balmus et al., 2018).

Therefore, to investigate the effects of reducing NAT10 function in tauopathy model animals, we generated heterozygous dNAT10 mutants expressing R406W MAPT (tau). As observed after Remodelin treatment, dNAT10 heterozygous null animals expressing MAPT R406W had fewer neuronal nuclear lamina invaginations than MAPT-R406W animals (Figure 6A and 6B). Furthermore, MAPT-R406W; heterozygous dNAT10 mutant animals had increased lifespan compared with MAPT-R406W alone (Figure 6C). Overall, reducing the activity of NAT10 *in vivo* corrects neuronal nuclear lamina defects and extends lifespan in a *Drosophila* model, consistent with the findings in human FTD-MAPT neurons.

**Figure 6.**
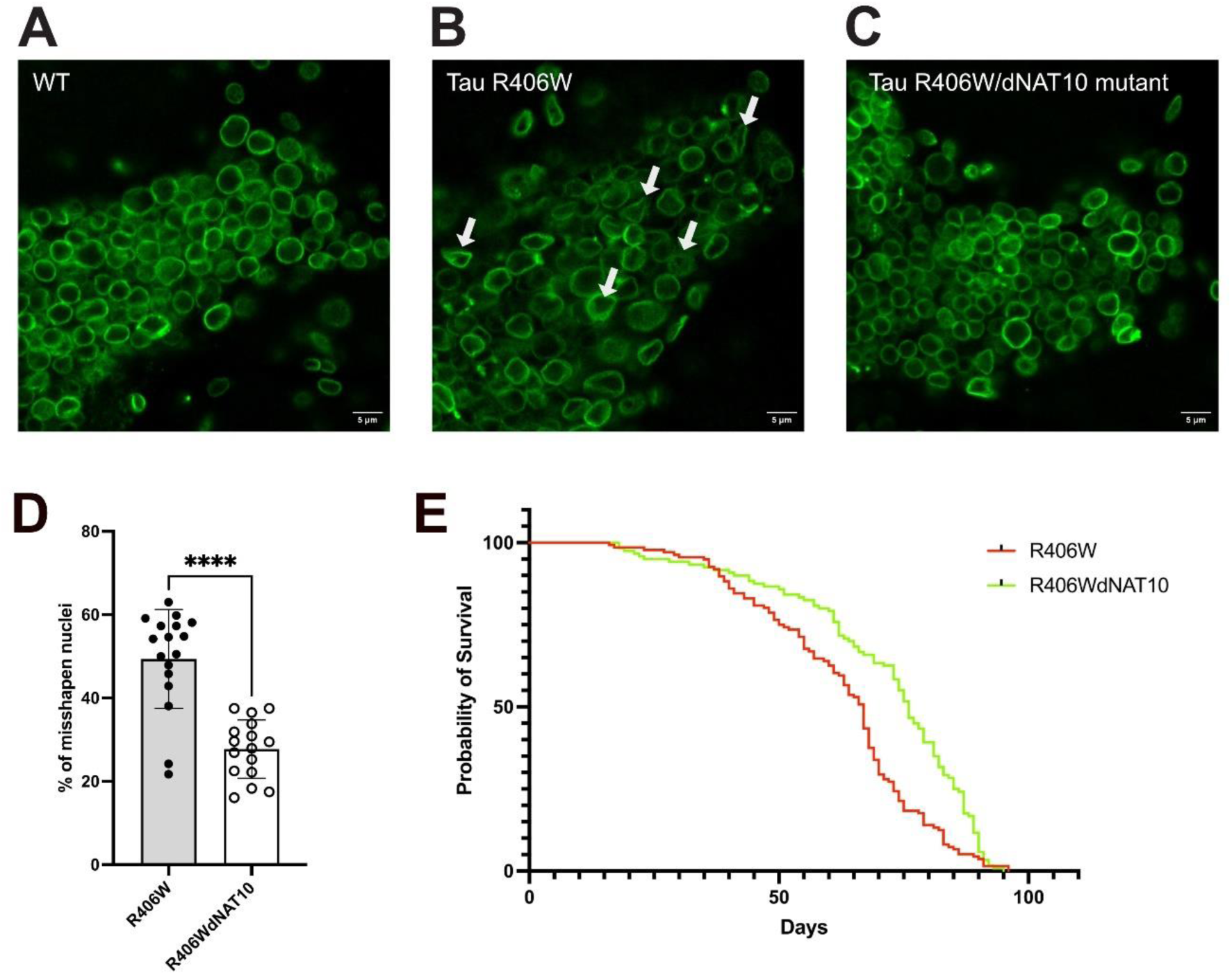
Haploinsufficiency of Drosophila NAT10 corrects nuclear lamina defects and extends lifespan in the MAPT R406W Drosophila tauopathy model. **A-C.** Confocal images of the nuclear lamina (Lamin B, green) of antennal lobe neurons in 20-day-old elaV > R406W *Drosophila* neurons compared to dNAT10 (+/-) elaV > R406W and control (w1118) neurons. Neurons in the antennal lobe displayed frequent nuclear lamina defects in MAPT R406W flies (arrows), whereas neurons in R406W, heterozygous dNAT10 mutant flies animals show improved nuclear morphology. Scale bars, 5 μm **D.** Quantification of misshapen nuclei was performed on 20-day-old *Drosophila* elaV > R406W neurons. Animals heterozygous for a mutant dNAT10 allele show a significantly reduced frequency of misshapen nuclei when compared to wildtype dNAT10 flies. Significance was tested using unpaired Student’s t test (p**** < 0.0001); error bar represents SD; n = 17 (R406W) and n = 16 (dNAT10(+/-) unique sections per brain hemisphere (max. 2 sections per animal), whereas each dot represents the percentage of misshapen nuclei per brain section scored. **E.** A longevity assay was performed to assess whether one copy of a mutant dNAT10 allele impacts lifespan in an R406W tauopathy model. dNAT10 (+/-) animals showed a significantly improved median lifespan. Significance was determined using the log-rank test (p****<0.0001); Survival was recorded for n = 135 and n = 120 for R406W and R406WdNAT10(+/-) female flies, respectively.

## DISCUSSION

We report here that the acetyltransferase NAT10 is a mediator of the pathogenic effect of tau in human frontotemporal dementia neurons *in vitro* and in a *Drosophila* tauopathy model *in vivo*. In both human FTD-MAPT and *Drosophila* MAPT R406W neurons, nuclear lamina defects are reversed by small molecule inhibition of NAT10 and by reduction of NAT10 protein levels. Correction of nuclear lamina defects in human FTD neurons also reversed defective nucleocytoplasmic transport, whereas NAT10 inhibition in a *Drosophila* tauopathy model reduced the neurotoxic effects of human mutant tau expression and extended lifespan.

Previously, we demonstrated that tau mutations causal for FTD led to tau accumulation in the neuronal cell body, nuclear lamina defects and nucleocytoplasmic transport dysfunction (Paonessa et al., 2019). Nocodazole-induced microtubule depolymerization acutely corrected nuclear lamina deformation, as we also observed in Hutchinson-Gilford Progeria (HGPS) fibroblasts (Larrieu et al., 2014). We previously reported that treatment with a small molecule inhibitor of NAT10, Remodelin, corrects nuclear lamina defects in HGPS, at least in part by releasing microtubule forces exerted on the nucleus (Larrieu et al., 2014). Given the similarities in nuclear phenotypes between HGPS and FTD-MAPT, we first investigated whether inhibition of NAT10 with Remodelin could reverse phenotypes in FTD-MAPT neurons.

We report here that NAT10 inhibition corrected nuclear lamina defects in human FTD-MAPT neurons. We extended these findings to an *in vivo* model of tauopathy, demonstrating that NAT10 inhibition and haploinsufficiency both significantly ameliorate nuclear shape abnormalities in a *Drosophila* model of tauopathy, MAPT R406W, where nuclear envelope defects have been previously reported (Frost et al., 2016). Highlighting the functional relevance of correcting tau-mediated pathologies, NAT10 inhibition and haploinsufficiency extended the lifespan of tauopathy model animals significantly.

Consistent with NAT10 inhibition having acute effects on neuronal cell biology, microtubule elongation was reduced in neurons within 24 hours of drug treatment. Furthermore, microtubule dynamics were only altered by NAT10 inhibition in FTD-MAPT neurons, with no effect on non-demented and isogenic control neurons. The widely accepted canonical function of tau protein is to stabilize axonal microtubules, however tau can also promote microtubule polymerization (Drubin and Kirschner, 1986). The findings presented here suggest that accumulation of tau in FTD-MAPT neurons results in a gain-of-function effect on microtubule dynamics, promoting microtubule formation in the neuronal cell body, altering their elongation and enabling transfer of mechanical forces to the nuclear membrane, deforming the nuclear lamina. Abnormal neuronal cell body microtubule dynamics, promoted by tau accumulation, may disrupt the nuclear membrane directly via LINC complex-mediated microtubule-nuclear envelope interactions (Kalukula et al., 2022). However, interactions between tau protein aggregates and the nuclear pore complex have been detected both in vitro and in vivo (Eftekharzadeh et al., 2018). Therefore, another possible mechanism for tau’s disruption of the nuclear membrane is via tau-mediated coupling of microtubules to the nuclear pore complex.

How NAT10 inhibition and loss of function correct microtubule-mediated nuclear membrane disruption in FTD-MAPT neurons is currently unclear. The acetyltransferase NAT10 is best characterized as a cytidine acetyltransferase that modifies several different forms of RNA, including ribosomal and tRNAs and protein-coding mRNAs (Achour and Oberdoerffer, 2024). However, NAT10 also acetylates a number of proteins, including histones, p53 and alpha-tubulin (Dalhat et al., 2021; Liu et al., 2016; Shen et al., 2009). We have previously confirmed that can NAT10 acetylate tubulin, and that this can be inhibited by Remodelin (Larrieu et al., 2014). Thus NAT10 may directly enhance microtubule stability via tubulin acetylation, a modification that promotes microtubule flexibility and overall stability (Ed et al., 2023). It is possible that microtubule acetylation in the cell body is reduced by NAT10 inhibition, combined with microtubule deacetylation by one or more of the known microtubule deacetylases, including HDAC6 and SIRT2 (Naren et al., 2023). In this case, reduced microtubule flexibility and stability would relieve the deforming forces on the nuclear envelope, as we have previously observed with acute microtubule depolymerization with nocodazole (Paonessa et al., 2019).

Overall, this study highlights the importance of nuclear lamina integrity and microtubule dynamics in the pathology of FTD and other tauopathies, and it proposes a novel therapeutic approach to target these cellular processes. Further research is warranted to fully understand the mechanisms by which Remodelin exerts its effects and to explore its potential as a treatment for neurodegenerative diseases characterized by tau pathology. Future studies should explore the long-term effects of Remodelin treatment *in vivo* and investigate whether these cellular improvements translate into functional benefits and neuroprotection in other animal models of tauopathies.

## METHODS

### Human iPSC lines

MAPT IVS10+16-A and MAPT IVS10+16-B mutant iPSCs were as reported in (Sposito et al., 2015). The non-demented control line was previously reported (Israel et al., 2012). The isogenic MAPT IVS10+16-B line was obtained through CRISPR/Cas9 editing of the MAPT IVS10+16-B line, reverting the mutant allele to the wild-type allele. iPSC cells were grown and expanded in feeder-free conditions using Essential 8 Medium (Thermo Fisher Scientific), at 37C with 5% CO2. Essential 8 Medium was replaced daily, and cells where routinely passaged using 0.5 mM EDTA when 60-80% confluent.

### Generation of iPSC-derived cortical neurons and drug treatments

Differentiation of iPSCs to cortical neurons was carried out as described, with minor modifications (Shi et al., 2012a, 2012b). Differentiated neurons were maintained in culture for up to 120 days. To establish identity and quality of cortical neuronal inductions, gene expression profiling was performed on a custom gene expression panel of approximately 250 genes (Strano et al., 2020). After subtracting the maximum negative control probe counts, gene counts were normalized using the geometric mean of 6 positive control probes and 7 housekeeping genes (CLTC, GAPDH, GUSB, PPIA, RPLP1, RPS15A, RPS9). For Remodelin (Sigma) treatment, neurons were grown for 120 days in vitro (DIV) and compound was added at 7.5 μM before imaging. DMSO was used as vehicle.

### Protein extraction and western blot analysis

Total cell protein was extracted with RIPA buffer (Sigma) supplemented with protease inhibitors (Sigma) and Halt phosphatase inhibitors (Thermo Fisher Scientific). Samples were run on NuPAGE 4%–12% Bis-Tris gels (Thermo Fisher Scientific) and proteins transferred to PVDF membrane (Millipore). All primary antibodies were incubated overnight in 5% milk in PBST at 4C. For detection, membranes were incubated for at least 1 h in secondary antibody, washed in PBST buffer and imaged on a LI-COR Odyssey CLx Infrared Imaging System.

### Immunofluorescence and imaging – human neurons

2D cultured cells on optically appropriate tissue culture plates were washed twice in PBS prior to incubation with PFA fixing solution for 15 minutes at room temperature. Cells were washed, incubated in blocking medium (5% v/v solution of normal donkey serum in TBS with 0.3% Triton-X) before primary antibody addition and incubation overnight at 4°C. Following primary antibody labelling, secondary antibodies were added in the presence of DAPI (SigmaAldrich, D9542). Immunostained samples were kept in TBS and stored at 4°C prior to imaging.

### Live imaging of microtubule dynamics

Neurons were grown to 100 DIV in individual m-Dish 35 mm dishes (Ibidi) and transfected with a plasmid encoding for GFP-EB3 (gift from Michael Davidson; Addgene plasmid # 56474). 48h after transfection, neurons were subjected to live imaging using a Leica SP5 microscope equipped with a controlled environment chamber (37C; 5% CO2). Images were acquired at resonant scanning with a 63x objective (1frame/sec). Resulting movies were analyzed using the plusTipTracker software (Applegate et al., 2011).

### Nucleocytoplasmic transport assay

Nucleocytoplasmic trafficking was analyzed by infection of 120 DIV human iPSC-derived neurons with the lentiviral CMV–NLS–tdTomato–NES construct. Two days post infection, iPSC neurons were subjected to FRAP analysis using a Leica SP8 inverted confocal microscope equipped with a live imaging chamber (Okolab). Signal from tdTomato was recorded every 0.3 second for 15 second before being bleached in the nuclear area, for 30 iterations of 40–60% laser power. Recovery was monitored every 0.3 second for 150 seconds. Recovery was normalized to the average of the pre-bleached signals. Results were elaborated using the Leica FRAP-wizard tool. Lentiviral-S-tdTomato was a gift from Jeffrey Rothstein (Addgene plasmid # 112579 ; http://n2t.net/addgene:112579 ; RRID:Addgene_112579)

### MAPT R406W *Drosophila* model

*Drosophila* carrying UAS-MAPT 0N4R R406W (kindly donated by Mel Feany) were crossed to w1118 ; + ; elaV-GAL4 (Bloomington Stock Center #8760) for pan-neuronal expression of MAPT R406W (Frost et al., 2016, 2014). w1118 animals were used as controls. l(1)G00020 mutant animals (Bloomington Stock Center #11474; dNAT10 with P{lacW} insertion at the transcriptional start site) were crossed to generate the l(1)G0020/FM7c ; + ; elaV-GAL4 line.

### Food Preparation

Fresh, liquid standard cornmeal-agar medium was kept in a water bath at 55°C. 4mL of liquid medium was transferred into vials and either 16uL vehicle (20% DMSO; 65% 2-hydroxypropyl-β-cyclodextrin solution, H5784 Sigma Aldrich; 15% Tween 80, P8074 Sigma Aldrich;(Balmus et al., 2018) or 16 μl Remodelin 25mM stock solution (100 μM final Remodelin food concentration) was added to the liquid cornmeal-agar food, thoroughly mixed and stored overnight in a dark chamber at 4°C before bringing to room temperature. Remodelin food ingestion was confirmed by red dye food staining (Supplementary Figure 2).

### Nuclear shape scoring - Drosophila

Nuclear shape was scored based on Lamin B (Developmental Studies Hybridoma Bank, ADL67.10) immunostaining. Nuclear morphology was scored according to features described (Janssen et al., 2022). Confocal imaging of 4% PFA-fixed brain samples was performed using a Leica SP8 confocal microscope with a 63x oil-immersion objective. Brains were imaged on the ventral side of the antennal lobe at 7uM depth, whereas max. one section was sampled and scored per brain hemisphere. Images were analyzed and scored with the open-source software Fiji (Schindelin et al., 2012).

### Lifespan

Lifespan assays were performed according to the protocol described (Piper and Partridge, 2016). 5 replicates of equal-sized groups of animals per condition and genotype (25-35 females) were transferred to food vials and maintained at 25°C in a 12:12 light-dark humidity-controlled incubator. Animals were transferred to fresh food vials every 3-4 days and dead animals were scored every 1-2 days.

### Statistics - Drosophila

Unless otherwise specified, data are presented as mean values of the number of independently conducted experiments indicated in the legend of each figure. Error bars represent the standard deviation (SD). Statistical analysis was performed using the Prism 10 analytical software (GraphPad). Unpaired Student’s t test was used to compare differences between two groups, assuming the data were normally distributed. One-way ANOVA followed by Tukey’s correction for multiple testing was used to analyze the differences between more than two groups. *** p < 0.001, **p < 0.01, *p < 0.05.

