## Supplementary Figures and Legends for "Targeting the acetyltransferase NAT10 corrects pathologies in human frontotemporal dementia neurons and extends lifespan in an in vivo *Drosophila* tauopathy model"

### SUPPLEMENTARY FIGURES AND MOVIES

**Supplementary Figure 1: Determination of the maximum tolerable, non-toxic dose of Remodelin for *Drosophila***

Dose-dependent developmental toxicity of Remodelin in *Drosophila* was assessed by addition of a range of volumes of a 25mM stock solution of Remodelin to liquid cornmeal agar. Developmental toxicity was scored as larval lethal, pupal lethal and eclosed died, versus eclosed viable. 100 µM Remodelin (16 ml of 25 mM stock solution; red box) was the highest concentration tolerated without an increase in developmental toxicity.

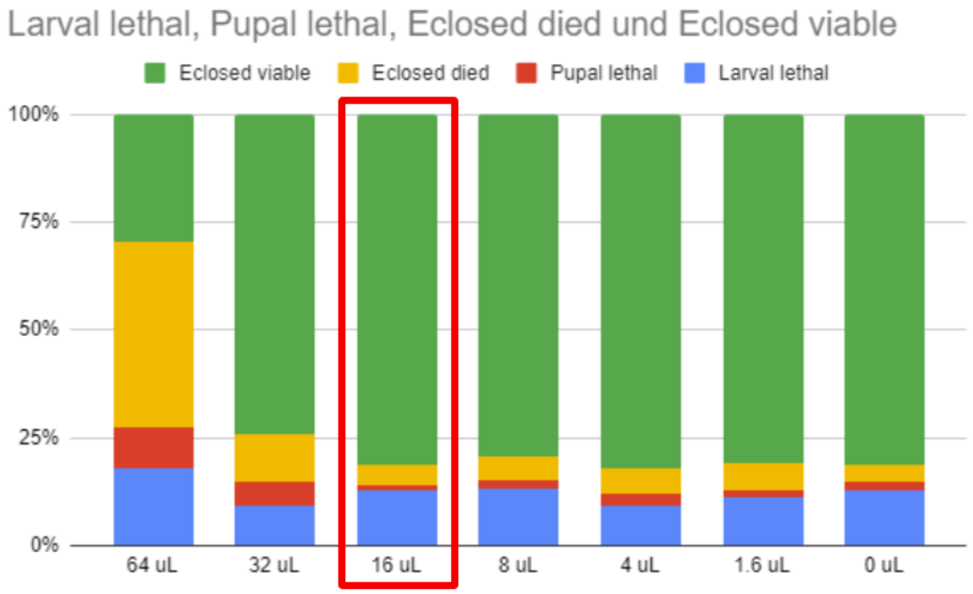

**Supplementary Figure 2: Confirmation of Remodelin ingestion by *Drosophila***

Non-toxic red food dye was added to the Remodelin-containing cornmeal agar food on which *Drosophila* were maintained. Ingestion of Remodelin was confirmed by the presence of red dye within the digestive tract/abdomen, as shown in the stereomicrograph below.

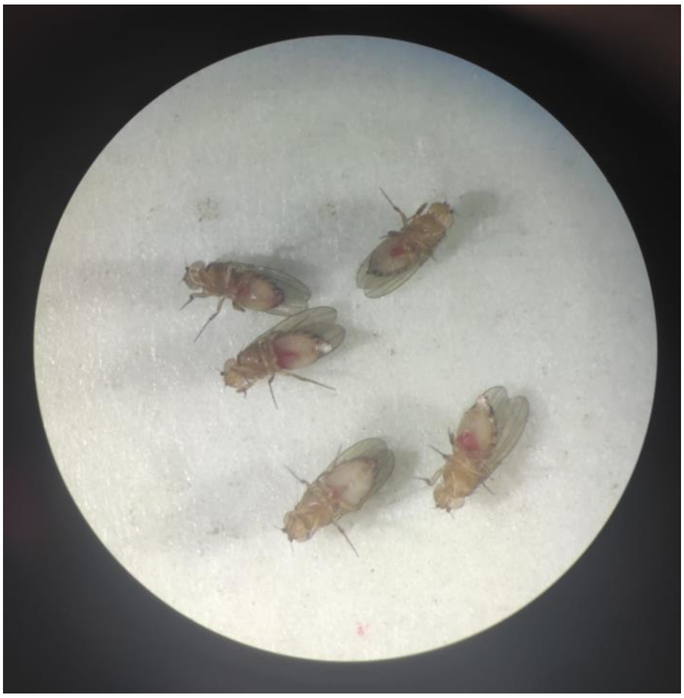

**Supplementary Movie 1: Live imaging of EB3-GFP in iPSC-derived non-demented control cortical neurons**

Confocal imaging of non-demented control cortical neurons expressing EB3-GFP treated with vehicle (DMSO, left) or Remodelin (right). Images were recorded for 150s, playback is accelerated to facilitate visualisation of microtubule movements.

**Supplementary Movie 2: Live imaging of EB3-GFP in iPSC-derived isogenic control cortical neurons**

Confocal imaging of non-demented control cortical neurons expressing EB3-GFP treated with vehicle (DMSO, left) or Remodelin (right). Images were recorded for 150s, playback is accelerated to facilitate visualisation of microtubule movements.

**Supplementary Movie 3: Live imaging of EB3-GFP in iPSC-derived cortical neurons from MAPT IVS10+16 patient 1**

Confocal imaging of non-demented control cortical neurons expressing EB3-GFP treated with vehicle (DMSO, left) or Remodelin (right). Images were recorded for 150s, playback is accelerated to facilitate visualisation of microtubule movements.

**Supplementary Movie 4: Live imaging of EB3-GFP in iPSC-derived cortical neurons from MAPT IVS10+16 patient 2**

Confocal imaging of non-demented control cortical neurons expressing EB3-GFP treated with vehicle (DMSO, left) or Remodelin (right). Images were recorded for 150s, playback is accelerated to facilitate visualisation of microtubule movements.
